# How a Predictive State Observer Can Self-Adapt Its Sensory Prediction-Error Correction Gain: Closed-Loop Evidence from a Muscle-Driven Reaching Task

**DOI:** 10.64898/2026.06.03.729790

**Authors:** Jun Kobayashi

## Abstract

We ask how a forward-model-based predictive state observer should set its sensory prediction-error correction gain during muscle-driven reaching, and whether that gain can be adapted from agent-available signals — innovation history and per-episode reaching outcome — rather than from swept oracle labels. We evaluate a residual-MLP forward model in a 34-muscle MyoSuite arm on an IK-reachable below-shoulder task, deployed in closed loop with a *stabilized endpoint probe controller* that uses non-negative least-squares muscle routing and a virtual target ramp; the controller is a stabilized probe for evaluating state-estimation effects, not a biological motor planner. A swept fixed-gain closed-loop oracle reveals a *delay-dependent* correction structure: with no sensory delay, intermediate correction gains are best (***K* = 0.25**–**0.50**), whereas with **18**-step delay observation-heavy correction wins (***K* = 1.0**). The forward-model-only ***K* = 0** ablation is not the oracle: it is systematically worse than the best fixed ***K*** by **1.9**–**6.1** cm and shows large NNLS controller residuals caused by long-horizon autoregressive drift; we therefore report ***K* = 0** as a diagnostic. Outcome-trained reliability-adaptive observers improve the delayed regime by **1.9**–**2.5** cm over default reliability while remaining neutral in no-delay cells, where the oracle is already intermediate. A feature-conditioned ***β*** adapter that maps cell-level innovation statistics to per-field gain parameters nearly matches a per-cell trained diagnostic in **5/6** cells, but both remain **1.4**–**1.8** cm worse than the swept fixed-***K*** oracle at **18**-step delay. These results separate the delay-dependent correction structure, the forward-model-only failure mode of ***K* = 0**, and the remaining limits of agent-available adaptive correction.

## 1 Introduction

State estimation in muscle-driven manipulation faces three coupled challenges: sensorimotor delay, observation noise, and signal-dependent motor noise. The internal-model view of biological motor control answers these with a *forward-model-based predictive state observer*: an internal forward model uses efference copy to predict the next state, and the observer corrects that prediction by a scalar weight *K* ∈ [0, 1] on the sensory prediction error Bernstein (1967); Wolpert and Ghahramani (2000). *K*=0 trusts the forward prediction (efference copy); *K*=1 trusts the sensor; intermediate *K* implements a sensory-prediction-error correction. The innovation-update form is structurally familiar from Kalman filtering, where the gain is derived from propagated covariance estimates Kalman (1960); Mehra (1970); Anderson and Moore (1979); we do *not* propagate covariances and do not claim to implement a full Kalman filter (Sec. 3.3). Instead we treat *K* as a scalar (or field-wise vector) that controls how strongly sensory prediction error corrects the current state estimate, and ask: *can the agent learn to set this gain from signals it generates online — innovation history and per-episode task outcome — rather than from per-condition K-sweep oracle labels?*

This question is biologically motivated. The nervous system does not have access to a swept oracle *K*^⋆^ across noise and delay regimes; it has access to its own prediction errors and, after moving, to whether the limb arrived where intended. We therefore propose a *two-layer reliability-adaptive observer. Within-trial*, a per-channel exponentially-weighted variance of squared innovation gives a running reliability estimate *r*_*f*_ (*t*), and a logistic *K*_*f*_ (*t*) = *σ*(*β*_0,*f*_ + *β*_1,*f*_ log *r*_*f*_ (*t*)) maps it to a field-wise correction gain on each of the five sensory fields into which we partition the 83-dimensional state—joint position and velocity, muscle activation, fingertip position, and reach_error (Sec. 3.4). *Across-trial*, Simultaneous Perturbation Stochastic Approximation (SPSA) Spall (1992) updates the meta-parameters *β* = {*β*_0,*f*_, *β*_1,*f*_} from the per-episode reaching outcome. Neither layer accesses the optimal *K*, the true state, or a swept oracle; both use only signals the agent generates during reaching. The resulting object is a computational internal-model loop in the cybernetic sense: efference-copy-based prediction is corrected by delayed sensory prediction error, while trial-level outcome reshapes the reliability-to-gain rule.

We instantiate this observer on a 34-muscle MyoSuite reaching arm Caggiano et al. (2022) running in the MuJoCo simulator Todorov et al. (2012) with a residual-MLP forward model trained under *H*-step rollout supervision (*H*=8) and a fixed-lag predictive state observer that handles configurable observation delay (*d* ∈ {0, 18} steps) and configurable per-field Gaussian noise. To deploy the observer in closed loop without confounding state-estimation effects with controller failure modes, we use a *stabilized endpoint probe controller*: a task-space PD on the tip error projected to non-negative muscle activations via non-negative least-squares (NNLS) routing, with a virtual target ramp that prevents impulsive endpoint commands at episode onset. The controller is a stabilized probe for evaluating state-estimation effects, not a model of biological motor planning.

The main task analyzed in this paper is restricted to IK-reachable targets below the neutral shoulder height (*z <* 1.393 m), giving a tabletop-like reaching distribution that avoids mechanically degenerate above-shoulder IK edge solutions.

Our contributions are summarized in four claims.

- **C1**. *The swept fixed-K closed-loop oracle is delay-dependent*. A 5-point *K*-sweep (*K* ∈ {0, 0.25, 0.5, 0.75, 1}) across the focused 3 noise × 2 delay grid, *n*=200 episodes per cell, reveals a delay-dependent correction structure: at delay-0 cells the best *K* is intermediate (*K* = 0.25–0.50, depending on noise level), while at delay-18 cells the best *K* is *K* = 1.0 across all noise levels (Fig. 2). The closed-loop oracle is therefore not uniformly prediction-heavy.
- **C2**. *K* = 0 *is a forward-model-only diagnostic, not the task oracle. K* = 0 is systematically worse than the best fixed *K* by 1.9–2.2 cm at delay-0 cells and 5.2–6.1 cm at delay-18 cells (paired 95% bootstrap CIs exclude zero in every cell; Fig. 3A). NNLS controller residuals at *K* = 0 are 92–96 across the grid, versus 11–49 for *K >* 0 (Fig. 3B), reflecting long-horizon autoregressive drift that the controller cannot realize as non-negative muscle activations. We therefore report *K* = 0 as a diagnostic ablation, not as an oracle.
- **C3**. *Outcome-trained reliability adaptation improves the delayed regime*. Across-trial SPSA on a single global *β* improves min-tip error over the default (*β*_0,*f*_ = 0, *β*_1,*f*_ = 0.5) rule by 1.85–2.46 cm at every delay-18 cell (paired 95% bootstrap CIs all exclude zero), while delay-0 cells are statistically neutral (Fig. 4). A single-cell SPSA run on a representative delayed cell achieves a 2.50 cm reduction over default on that cell.
- **C4**. *Feature-conditioned β adaptation nearly matches the per-cell-trained diagnostic for most cells, but both remain below the swept fixed-K oracle*. A 30-parameter linear adapter that maps z-scored log innovation statistics to per-field *β* matches a per-cell SPSA diagnostic in 5*/*6 cells (paired CIs cover zero); per-cell wins in one zero-delay high-noise cell only, by 1.2 cm (Fig. 4). However, both feature-conditioned *β* and per-cell *β* remain 1.4–1.8 cm below the swept fixed-*K* oracle at every delay-18 cell (paired CIs all exclude zero). Field-wise realized *K* confirms that the adaptive rule increases the joint-position gain from 0.72–0.92 (default) to 0.87–0.98 (SPSA) and suppresses the joint-velocity gain from 0.39–0.43 to 0.34, but the residual gap to the swept oracle persists, indicating that parameterization, reliability features, and observer dynamics jointly bound the achievable adaptation (Fig. 5).

**Fig. 1.**
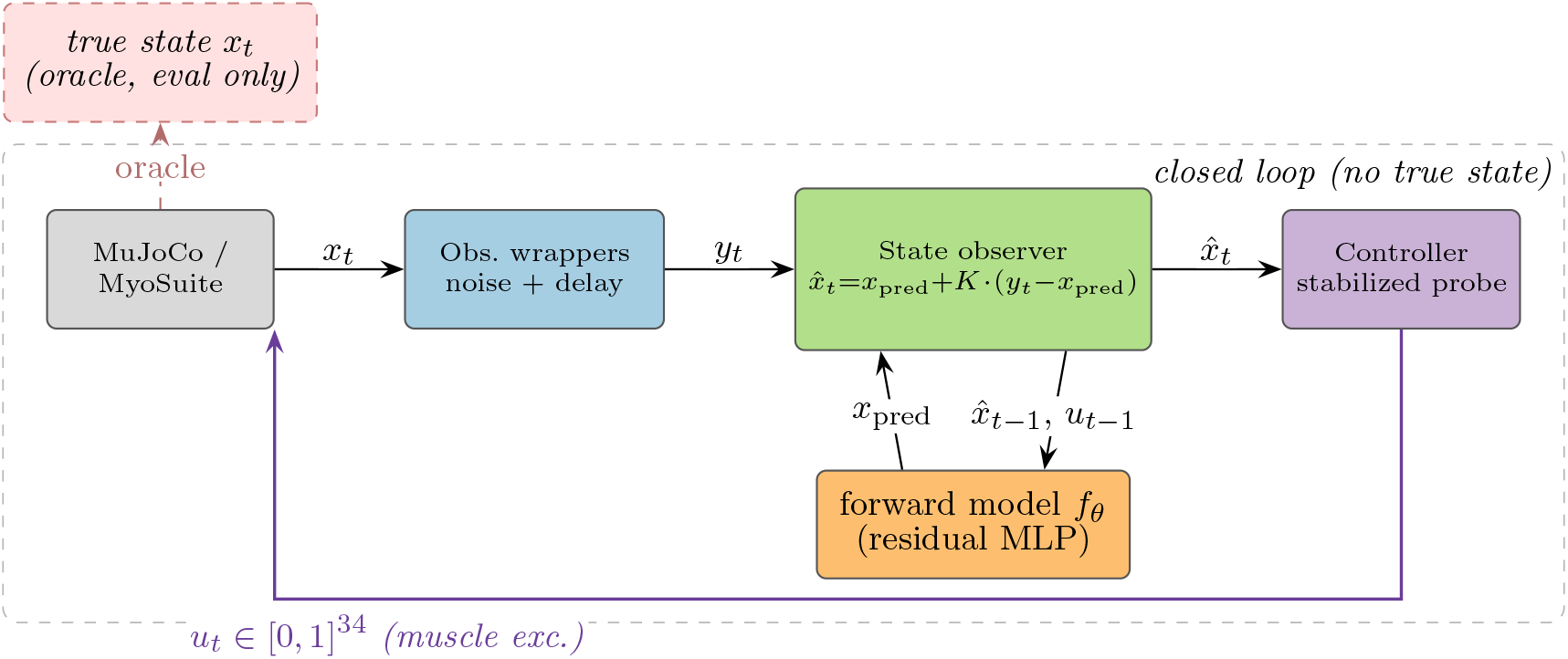
Closed-loop pipeline. Each control step, the environment’s true state *x*_*t*_ is filtered through observation wrappers that add per-field Gaussian noise and a fixed delay, producing *y*_*t*_. The predictive state observer combines *y*_*t*_ with a one-step prediction *x*_pred_ produced by the residual-MLP forward model 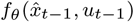 via a scalar correction gain *K* ∈[0, 1] on the sensory prediction error *y*_*t*_ − *x*_pred_, as in Eq. (4). The controller consumes only the observer output 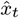 and emits the next muscle excitation *u*_*t*_ ∈ [0, 1]^34^. The true state *x*_*t*_ is exposed only to the offline evaluator (dashed red box); the closed loop never accesses it.

**Fig. 2.**
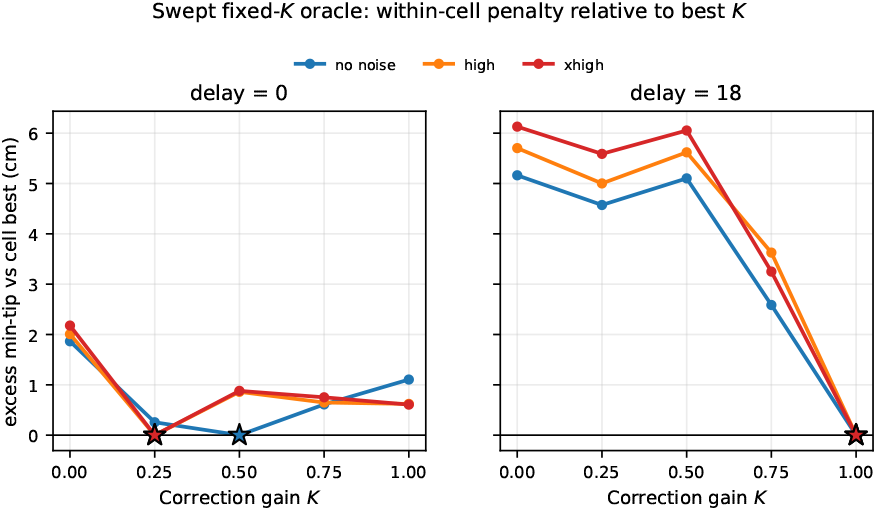
Swept fixed-*K* closed-loop oracle is delay-dependent. Excess min-tip error relative to the best fixed *K* within each noise/delay cell (*n*=200 episodes/cell) under the stabilized endpoint probe across *K* ∈ {0, 0.25, 0.5, 0.75, 1}, split by delay. Colored stars mark the per-cell best *K* at zero penalty. This visualization emphasizes the shape of the *K* curve within each cell rather than comparing absolute task difficulty between delays. Delay-0 cells (left) prefer intermediate *K*, shifting toward prediction as observation noise rises. Delay-18 cells (right) collapse to *K* = 1 uniformly. The closed-loop oracle is therefore delay-dependent rather than uniformly prediction-heavy.

**Fig. 3.**
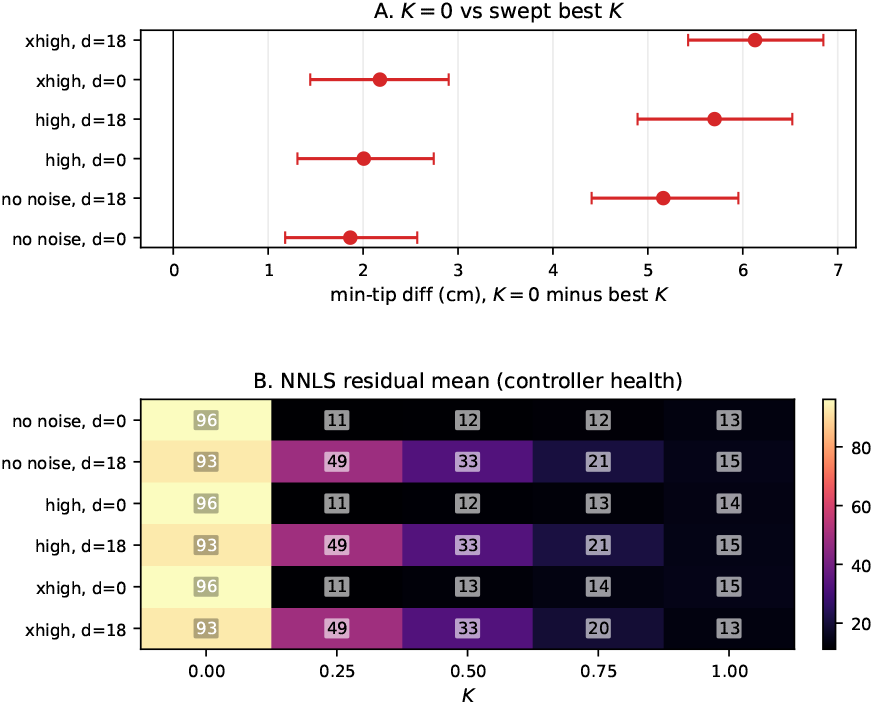
*K*=0 is a forward-model-only diagnostic, not the closed-loop oracle. (A) Per-cell paired min-tip difference *K*=0 minus best fixed *K*, in cm, with 95% percentile bootstrap CI (10,000 resamples, paired by episode index); *K*=0 is systematically worse in every cell. (B) NNLS residual mean ∥*M* ^⊤^*a* − *u*_joint_∥ _2_ as a heatmap over *K* and cell: *K*=0 runs sit far above the rest of the grid, indicating the requested joint command is unrealizable in non-negative muscle space.

**Fig. 4.**
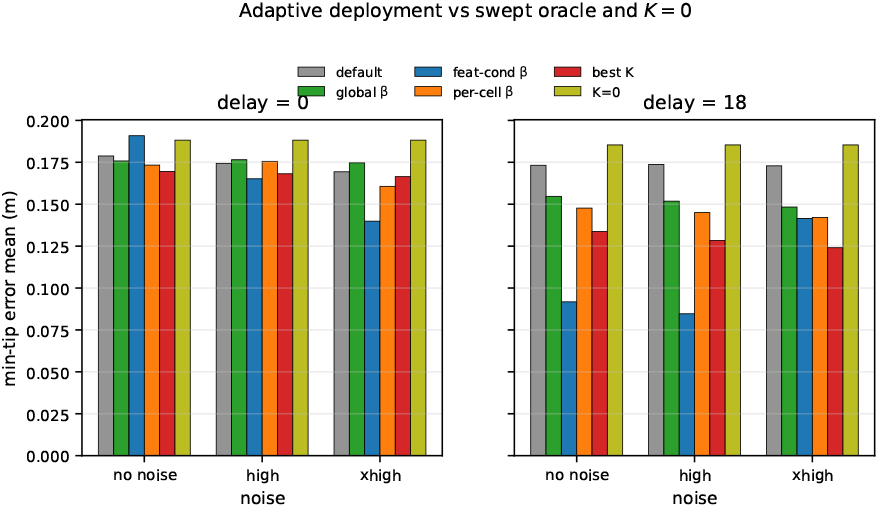
Outcome-trained reliability adaptation in closed loop. Per-cell min-tip error mean (*n*=200 episodes/cell) under the stabilized endpoint probe. Bars: default reliability (gray), global SPSA *β* (green), feature-conditioned *β* (blue), per-cell SPSA diagnostic (orange), per-cell best fixed *K* (red), *K*=0 diagnostic (yellow). Adaptive estimators improve over default at delay-18 cells but remain above the swept oracle by 1.4–1.8 cm. At delay-0 cells, adaptive estimators sit near default; the oracle there is intermediate *K* rather than observation-heavy.

**Fig. 5.**
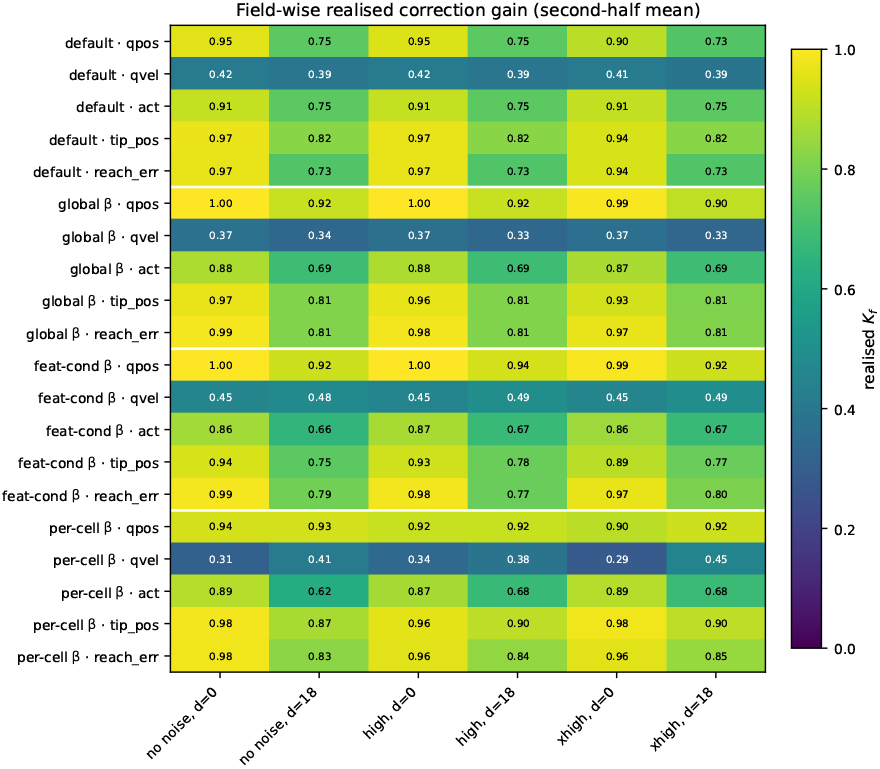
Field-wise realized correction gain (second-half episode mean) under the four deployed estimators. Rows: estimator × field (5 fields per estimator); columns: 6 cells. Default reliability (top block) sits at moderate gains across fields; global SPSA (second block) increases the `qpos` gain and suppresses `qvel`; feature-conditioned and per-cell adapters (lower blocks) shift field-wise trust per cell. The reliability-to-gain rule does adapt along field-wise directions, but the residual oracle gap of C4 is not eliminated.

Together, C1–C4 separate three things that an earlier controller diagnostic conflated: the swept-*K* oracle structure (delay-dependent, not uniformly *K* = 0); the *K* = 0 failure mode (forward-model-only diagnostic, not a usable oracle); and the remaining limits of agent-available adaptive correction (observation-heavy regimes can be reached, intermediate-*K* regimes cannot, even with cell-conditioned *β* or per-cell training).

The rest of the paper is organized as follows. Section 2 positions our work against prior literature on adaptive correction gains and Kalman-style updates, forward models in motor control, learned dynamics, outcome-driven adaptation, and muscle-driven simulation. Section 3 describes the below-shoulder reaching task, the forward model, the fixed-lag predictive state observer, the reliability-adaptive correction-gain rule, the across-trial SPSA loop, the stabilized endpoint probe controller, and the evaluation pipeline. Section 4 reports the four claims with the corresponding figures: the delay-dependent fixed-*K* oracle structure, the *K* = 0 diagnostic, outcome-trained reliability adaptation in delayed reaches, and the remaining gap to the swept oracle. Section 5 discusses the trade-offs and the implications of agent-available correction-gain adaptation for biological motor control models, the role of controller-health diagnostics in closed-loop estimator benchmarks, and the limits of forward-model-only deployment exposed by the *K* = 0 ablation.

## 2 Related Work

### Adaptive correction gains and Kalman-style updates

The Kalman filter Kalman (1960) and its smoothing counterpart Rauch et al. (1965) are the textbook framework for combining a state-space model with noisy observations under known noise covariances. When the covariances are unknown, the classical *adaptive Kalman filter* estimates them online from the innovation sequence Mehra (1970); the steady-state gain *K* then drops out of the covariance estimate. We adopt the same innovation-style update form (residual MLP for dynamics, fixed-lag buffered correction for delays) but *do not* propagate covariances and do not compute an optimal Kalman gain; the state-space formulation is standard text-book material Anderson and Moore (1979) and we treat the result as a predictive state observer with a scalar correction gain (Sec. 3.3). The novelty of our setting is therefore not in the filter equations but in the *choice* of *K*: instead of deriving *K* from a noise-covariance estimate, we first measure the closed-loop fixed-*K* oracle and then ask whether outcome-trained, agent-available reliability adaptation can approach it without seeing oracle labels. This closed-loop, task-outcome-driven framing is largely absent from the adaptive-Kalman literature and is not natural inside the covariance-propagation view.

### Forward and internal models in motor controll

The view that the central nervous system maintains an internal forward model of the limb to predict the sensory consequences of motor commands traces back to Bernstein’s degrees-of-freedom analysis Bernstein (1967) and was given an explicit computational framing by Wolpert and Ghahramani Wolpert and Ghahramani (2000), who identified forward and inverse models as complementary processes underlying sensorimotor estimation, prediction, and learning. Our residual MLP is a coarse instantiation of the forward model in that picture; the multi-step supervision we add is motivated by the fact that the loop that consumes the estimate operates over hundreds of integration steps and thus depends on long-horizon prediction quality. More biologically grounded forward-model families — continuous-time recurrent networks with input-modulated time constants such as liquid time-constant networks Hasani et al. (2021) and their closed-form variant Hasani et al. (2022) — have been proposed for sensorimotor dynamics; in this work we keep the forward-model implementation intentionally generic to isolate the oracle-choice question.

### Learned dynamics and multi-step supervision

Model-based reinforcement learning systems have moved from one-step dynamics losses to multi-step rollout objectives because closed-loop planners need predictions that survive unrolling. Dreamer Hafner et al. (2020) backpropagates through imagined trajectories in a latent world model, and MBPO Janner et al. (2019) branches short model rollouts from real data, formalizing “when to trust your model”. We take the same lesson into observer design: a forward model that is accurate enough over the controller’s planning horizon shifts where prediction-error correction is helpful and where it is harmful.

### Outcome-driven adaptation and SPSA

Outcome-driven, gradient-free adaptation has a long history in motor control: trial-by-trial visuomotor adaptation in humans Wolpert and Ghahramani (2000) is captured by an internal-model update rule that uses only the observed end-of-trial error, not the underlying optimal-gain calculation. Algorithmically, our across-trial loop follows Spall’s Simultaneous Perturbation Stochastic Approximation (SPSA) Spall (1992): a two-point gradient estimate using a single Rademacher perturbation per iteration, with a robust hyperparameter schedule. SPSA’s appeal in this setting is that it needs only the per-episode outcome, not the per-step state, and that its computational cost scales with the number of meta-parameters *β* (here, 10) rather than the dimension of the observer state (here, 83). Behavior cloning and DAgger Pomerleau (1988); Ross et al. (2011) were used in an earlier version of this work to ask whether the resulting controllers consume state estimates; we found that with small demonstration sets, they do not consistently expose observer differences, and we now use only state-coupled feedback controllers in the main results, so the observer signal flows to muscle commands.

### Muscle-driven simulation

Our task runs in MuJoCo Todorov et al. (2012) via the MyoSuite musculoskeletal benchmark Caggiano et al. (2022), which provides a 34-muscle reaching arm and a contact-rich set of tasks suitable for sensorimotor estimation studies. MyoSuite is increasingly used as a testbed for reinforcement learning controllers; here, we use it instead to ask whether a predictive state observer can adapt its correction gain from agent-available signals alone, a question that the RL-centric literature has not addressed.

## 3 Method

### 3.1 Task and Environment

We use the `myoArmReachFixed-v0` task in MyoSuite 2.12.2, a 34-muscle shoulder-elbow-wrist-hand system actuated by activations in [0, 1]^34^. The task is to drive the index-finger tip to a fixed target in a 12 s episode at Δ*t*=0.02 s (600 control steps). The flat state *x* ∈ ℝ^83^ stacks joint positions *q* ∈ ℝ^20^ (`qpos`), joint velocities 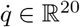 (`qvel`), muscle activations *a* ∈ ℝ^34^ (`act`), fingertip position *p*_tip_ ∈ ℝ^3^ (`tip_pos`), target *p*_tgt_ ∈ ℝ^3^ (`target_pos`), and reach_error *e*=*p*_tip_ − *p*_tgt_ ∈ ℝ^3^ (r`each_err`). True state is extracted from MuJoCo and used only as an oracle for evaluation; the closed loop never accesses it (Fig. 1).

#### Below-shoulder target set

Targets are sampled uniformly from the environment’s task-space reach range and accepted if a damped-Jacobian IK solver places the tip within 1 cm and the target lies below the neutral shoulder height (*z <* 1.393 m). The shoulder constraint avoids mechanically degenerate above-shoulder IK edge solutions and yields a tabletop-like reaching distribution; it is anatomically motivated rather than a purely numerical post-hoc filter. We generate *n*=200 accepted targets with seed offset 30,000; acceptance rate is 55.9% (rejections are due to the shoulder-height criterion, not IK failures). Appendix C summarizes the diagnostic rationale for this target restriction.

### 3.2 Forward Model

A residual MLP *f*_*θ*_(*x*_*t*_, *u*_*t*_) predicts the state delta Δ*x*_*t*_=*x*_*t*+1_ − *x*_*t*_ rather than *x*_*t*+1_ directly. At Δ*t*=0.02 s the state changes slowly, so *x*_*t*+1_ ≈ *x*_*t*_ component-wise; predicting Δ*x* frees the model from re-learning the identity map and concentrates capacity on the small per-step dynamics that the filter and controller actually consume. The parameterization is standard in model-based

RL world models Hafner et al. (2020); Janner et al. (2019). The input is the flat 83-dimensional state and the 34-dimensional excitation; the architecture is two hidden layers of 256 units with ReLU and LayerNorm; training uses Adam Kingma and Ba (2015). In the single-step baseline, *f*_*θ*_ is trained on the standard one-step MSE loss

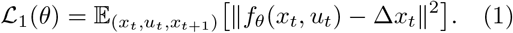

#### Multi-step rollout supervision

The multi-step variant trains *f*_*θ*_ with an *H*-step rollout loss instead of the one-step MSE, where *H* denotes the training rollout horizon. For each contiguous trajectory chunk (*x*_*t*_, *u*_*t*_, …, *x*_*t*+*H*_) inside a single source episode, the model is unrolled free-running: 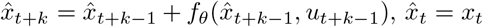, and the loss averages MSE across the *H* prediction steps:

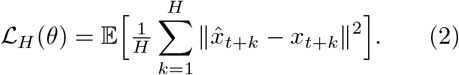

Multi-step rollout supervision has been used to stabilize long-horizon predictions in latent-imagination world models Hafner et al. (2020) and in model-based policy optimization Janner et al. (2019), where the relevant horizon is the planner’s rollout depth rather than the observer’s delay; we re-use the technique here to make *f*_*θ*_ robust across the observer’s buffer rollout.

We train the same architecture with *H* ∈ {1, 4, 8} on the expanded reachable-target dataset (77 k transitions across 150 episodes; see Appendix B).

### 3.3 Fixed-Lag Predictive State Observer

The observer uses an innovation-update form structurally familiar from Kalman filtering, but does not propagate covariances and does not compute an optimal Kalman gain. The gain *K* ∈ [0, 1] is a *sensory prediction-error correction gain*, not a covariance-derived Kalman gain.

The observation pipeline produces *y*_*t*_ = *x*_*t*−*d*_ + *η*_*t*_, where the observation shares the same schema as the 83-dimensional state vector, *d* ∈ ℤ_≥0_ is a fixed delay in control steps, and *η*_*t*_ is per-field i.i.d. Gaussian noise with diagonal covariance diag(*σ*^2^). The estimator maintains a length-(*d*+1) ring buffer of past one-step estimates 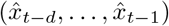 together with the corresponding applied actions (*u*_*t*−*d*_, …, *u*_*t*−1_). At each step, the observer first runs the forward model on every buffered estimate to obtain a prediction at the present time,

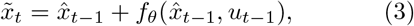

then forms an innovation against the *delayed* buffer entry and applies an element-wise gain *K* ∈ [0, 1]^83^:

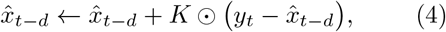

and finally re-rolls the corrected past estimate forward through the buffered actions to obtain the present-time estimate 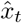. Eq. (4) collapses to the standard one-step innovation update when *d*=0. Throughout the paper *K* is a scalar broadcast across the 83 state coordinates so the correction gain reduces to a single trust knob between forward prediction (*K*=0) and observation (*K*=1). This fixed-lag construction follows the buffered Rauch–Tung–Striebel form Rauch et al. (1965); Anderson and Moore (1979) specialized to a scalar gain.

#### Noise and delay grids

The main closed-loop experiments use a focused grid with delays *d* ∈ {0, 18} and three observation-noise presets. We denote the categorical noise level by *ν* ∈ {none, high, xhigh}. The label *none* sets all observation-noise standard deviations to zero; *high* uses *σ*_qpos_ = *σ*_qvel_ = 0.02 and *σ*_tip_ = *σ*_reach_ = 0.01; *xhigh* uses *σ*_qpos_ = *σ*_qvel_ = 0.08 and *σ*_tip_ = *σ*_reach_ = 0.04. These labels are used only as compact table and figure notation for the corresponding per-field Gaussian noise settings (the sensory fields are defined in Sec. 3.4).

### 3.4 Reliability-Adaptive Correction Gain (Within-Trial)

The observer above uses a fixed scalar *K* broadcast across all 83 fields. We propose instead an *agent-available, field-wise reliability-adaptive* rule that sets *K*_*f*_ (*t*) online from each sensory channel’s own innovation history. No oracle on the optimal gain, no true state, and no swept-condition labels enter the rule; only the per-step innovation 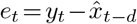 does.

We partition the 83-dimensional state into five adaptive *sensory fields*—20-D `qpos`, 20-D `qvel`, 34-D `act`, 3-D `tip_pos`, and 3-D `reach_err—` and a sixth, 3-D fixed-gain target channel, `target_pos`. The partition is an engineering grouping driven by the MyoSuite state schema, but it admits a coarse biological reading: `qpos` and `qvel` are proprioceptive-like, `act` is efferent and muscle-command-like, and `tip_pos` and `reach_err` are visual and task-space-like. The 3-D target-position channel is held at a fixed gain because the target is known and does not have a meaningful prediction error. For each field *f ∈ ℱ*= {qpos, qvel, act, tip pos, reach_err} the observer maintains an exponentially-weighted (EMA) estimate of the per-field squared-innovation mean,

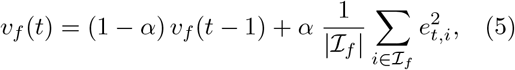

where ℐ_*f*_ are the field’s coordinate indices in the 83-dimensional state vector and *α* = 0.05 unless otherwise noted. The agent’s running estimate of *sensory reliability* for field *f* is the inverse of this variance,

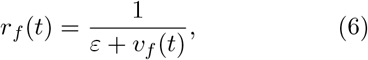

with *ε* = 10^−6^. We then map reliability to a per-field correction gain through a two-parameter logistic,

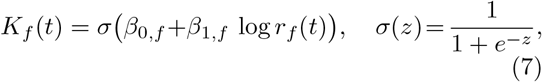

and form the 83-dimensional gain vector *K*(*t*) ∈ [0, 1]^83^ by broadcasting *K*_*f*_ ∈ (*t*) over its field’s coordinates. The innovation update in Eq. (4) is then applied elementwise with *K*(*t*) in place of the scalar *K*.

Eq. (7) is intentionally simple. The slope *β*_1,*f*_ controls how sharply the gain reacts to changes in log-reliability, and the intercept *β*_0,*f*_ controls the operating point at *r*_*f*_ (*t*) = 1 (i.e., *K*_*f*_ = *σ*(*β*_0,*f*_)). With *β*_0,*f*_ = 0, *β*_1,*f*_ = 0.5, and *v*_*f*_ (0) = 1, the rule produces *K*_*f*_ (*t*) ≈ 0.5 on the first step and adapts upward when innovation shrinks (the sensor is informative) or downward when innovation grows (the sensor is noisy). Per-field *β* = {*β*_0,*f*_, *β*_1,*f*_} lets the rule learn that different fields have different reliability–correction trade-offs under the same physical noise level.

### 3.5 Across-Trial Outcome Adaptation (SPSA)

The meta-parameters *β* = {(*β*_0,*f*_, *β*_1,*f*_): *f* ∈ℱ ∈ ℝ^10^ themselves can be adapted across trials from *outcome* signals the agent has at the end of each episode. We define the per-episode outcome as the minimum tip-to-target distance,

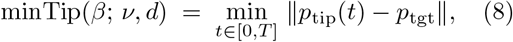

computed from the simulated trajectory after each episode. We use minTip as an *operational scalar outcome* for across-trial optimization, not as a biological claim that an agent directly observes the trajectory minimum. The across-trial loop minimizes 𝔼[minTip(*β*)] where the expectation is over target draws (and, in the multi-cell variant, over uniformly drawn (*ν, d*) cells from the focused grid).

Because the closed-loop minTip is not differentiable in *β* (it passes through environment steps, EMA state, and a non-smooth min), we use Simultaneous Perturbation Stochastic Approximation (SPSA) Spall (1992): at iteration *n*, draw a Rademacher vector Δ_*n*_ ∈{±1}^10^, evaluate the outcome at *β*_*n*_ ± *c*_*n*_Δ_*n*_ averaged over *S* paired episodes (samples per side), form the central-difference gradient estimate

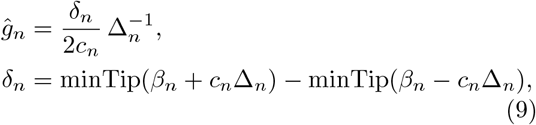

and update *β*_*n*+1_ = *β*_*n*_ − *a*_*n*_*ĝ*_*n*_. The Spall schedules are 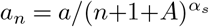 and 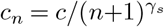 with *a*=2.0, *c*=0.3, *α*_*s*_=0.602, *γ*_*s*_=0.101, and *A*=5. Two SPSA variants are reported: *single-cell* (*S*=12, 100 iterations, no-noise, *d*=18) and *full-grid* (*S*=12, 100 iterations, uniformly drawn (*ν, d*) from the 6-cell grid per iteration).

### 3.6 Feature-Conditioned *β* Adapter

§3.5 optimizes one global *β* ∈ ℝ^10^ and deploys that same reliability-to-gain map in every noise/delay cell. The feature-conditioned adapter asks whether *β* can be adjusted from information available inside the cell itself: the magnitude and variance of each field’s recent innovation. We start from a fixed base parameter *β*^base^ and add a small independent linear correction for each sensory field.

#### Architecture

For each sensory field *f* ∈ ℱ, the adapter outputs a correction (Δ*β*_0,*f*_, Δ*β*_1,*f*_) from two per-field features:

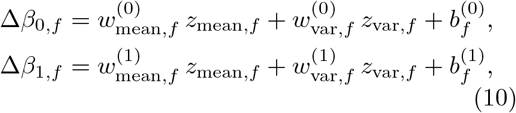

giving five fields × two outputs × three weights = 30 parameters total. The effective per-field *β* used by the within-trial layer is

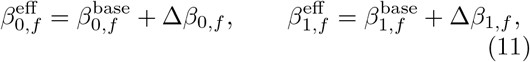

fixed for the duration of one episode. Setting all 30 adapter weights to zero recovers *β*^base^ exactly, so the adapter is an additive correction. Throughout the main results we take *β*^base^ to be the global SPSA *β* trained on the focused grid (Sec. 3.5).

#### Input features and normalization

The two per-field features *z*_mean,*f*_ and *z*_var,*f*_ are agent-available innovation statistics: mean absolute innovation and mean squared innovation over the second half of a brief default-reliability warm-up run on the cell. Each statistic is then log-transformed and z-scored *across the focused-grid cells* using normalization constants estimated once from the warm-up rollouts and then frozen:

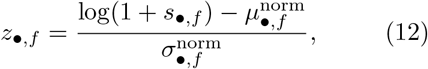

where *s*_•,*f*_ is the raw statistic and (*µ*^norm^, *σ*^norm^) are calibration constants. This calibration is analogous to sensory scale calibration in a biological agent: it adjusts the units of innovation features into a common scale, but does not encode the optimal correction gain or any oracle label.

#### Training

The 30 adapter weights are trained by SPSA on the same per-episode min-tip outcome used in Sec. 3.5, with 200 iterations, *S* = 12 paired episodes per iteration (two samples per cell, six cells), perturbation amplitude *c* = 0.3 and learning rate *a* = 0.5, both annealed by the Spall schedule. The cell features computed in the calibration step are reused at each iteration without modification — the adapter learns a single context-to-*β* map, not a per-cell *β*.

### 3.7 Controllers

All main closed-loop results use a single *stabilized endpoint probe controller*. We retain it only because we need a closed-loop probe that does not by itself fail in muscle space; it is not a model of biological motor planning and not a candidate optimal-control policy.

#### Stabilized endpoint probe

A task-level PD law on the estimated tip-to-target error is projected into the non-negative muscle-activation cone via non-negative least squares against the actuator moment-arm matrix *M* ∈ ℝ^34×20^, and the controller’s effective target is ramped from the initial tip position 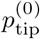 to the episode target *p*^⋆^ over *T*_ramp_=300 steps to avoid impulsive commands at episode onset. At control step *t*, with estimated tip position 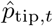 and estimated joint velocity 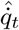,

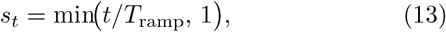

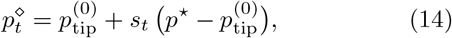

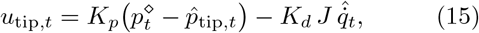

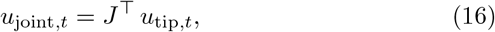

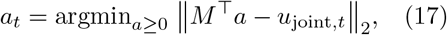

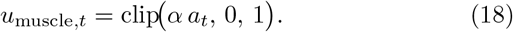

Here *s*_*t*_ is the ramp fraction, 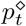 is the virtual target, *u*_tip,*t*_ and *u*_joint,*t*_ are task-space and joint-space demand vectors, *a*_*t*_ is the non-negative NNLS routing variable, and *u*_muscle,*t*_ is the final muscle command. We use *K*_*p*_=30, *K*_*d*_=3, *α*=5, with *J* the tip-site positional Jacobian captured at the environment’s neutral pose. NNLS is the SciPy implementation of the standard active-set NNLS algorithm.

The NNLS routing in Eq. (17) is used because it (i) respects the non-negative muscle constraint without ReLU-induced symmetry-breaking between antagonist pairs and (ii) returns a residual norm ∥*M* ^⊤^*a* − *u*_joint_∥ _2_ that we log as a per-step controller-health diagnostic. The virtual target ramp in Eqs. (13)–(14) is a transparent demand-shaping device, not a minimum-jerk planner; we deliberately do not couple it to an optimal-control objective.

We refer to this controller as a *stabilized probe controller* throughout. Appendix C summarizes the controller diagnostic that motivated this stabilized design.

### 3.8 Evaluation Pipeline

*Open-loop* evaluation replays a recorded episode and applies observation wrappers to synthesize *y*_*t*_ for an arbitrary (*ν, d*) combination; the estimator is then run on the synthesized stream and per-field state error is aggregated. *Closed-loop* evaluation drives the actual environment: *u*_*t*_ comes from the controller, which sees the estimator output (never the true state). The main closed-loop fixed-*K* sweep and adaptive deployments use 200 episodes per noise/delay cell with paired episode seeds across estimators. Smaller 10-episode sweeps are used only for historical diagnostics in Appendix C. The open-loop diagnostic grid uses 3 controllers × 6 noise levels × 4 delays; the main closed-loop grid uses the focused 3 noise levels × 2 delays defined above.

## 4 Results

### 4.1 Forward-model selection and h=600 long-horizon diagnostic

The main forward model is a residual MLP trained with *H*=8 rollout supervision on the 77,292-transition reachable-target dataset (Appendix B); it attains an *h*=50 tip prediction error of 0.034 m (full rollout metrics in Table B2), which meets the open-loop validation threshold, and is the model used in all closed-loop results below.

Because the closed-loop episode horizon is 600 control steps, we additionally run a zero-action *h*=600 autoregressive stability diagnostic on the four forward-model variants (*H*=1, *H*=4, *H*=8 main, undertrained *H*=8; Appendix B). The two *H*=8 variants keep |*q*| _max_ ≤ 5 rad and 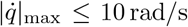 under zero action; for the main *H*=8 model, tip drift of 1.03 m is at the anatomical-workspace boundary. The state-vector *p*_tgt_ field drifts by 0.54 m even though the environment target is invariant, a known structural limitation of including the target field in the predicted state, which we leave as future work. We retain target drift as a diagnostic field, but do not include it in the formal open-loop validation threshold (Appendix B).

### 4.2 Fixed-*K* sweeps reveal delay-dependent correction structure (C1)

We run a 5-point closed-loop *K*-sweep (*K* ∈ {0, 0.25, 0.5, 0.75, 1}) under the stabilized end-point probe on the focused 3 noise × 2 delay grid, *n*=200 episodes per cell, with the below-shoulder target set (Sec. 3.1). Each estimator is a fixed-gain predictive state observer over the *H*=8 forward model.

Figure 2 plots the within-cell penalty as a function of *K*, defined as excess min-tip error relative to the best fixed *K* in the same noise/delay cell. This normalization is only for visualization; the absolute cell means are reported in Table 1. The correction structure is delay-dependent. At delay-0 cells, the best fixed *K* is intermediate: *K* = 0.50 for *ν*=none, *K* = 0.25 for *ν*=high and *ν*=xhigh. Increasing observation noise pushes the no-delay optimum from *K* = 0.50 to *K* = 0.25, i.e., the estimator favors its forward-model prediction more strongly when the channel is noisier — the qualitative direction expected from reliability-weighted prediction-error correction. At delay-18 cells, the best fixed *K* collapses to *K* = 1.0 across all three noise levels; the stale forward-model prediction (delayed by the same 18 steps as the observation) is no longer a useful current-state anchor, and observation-heavy correction wins despite the noise.

**Table 1.**
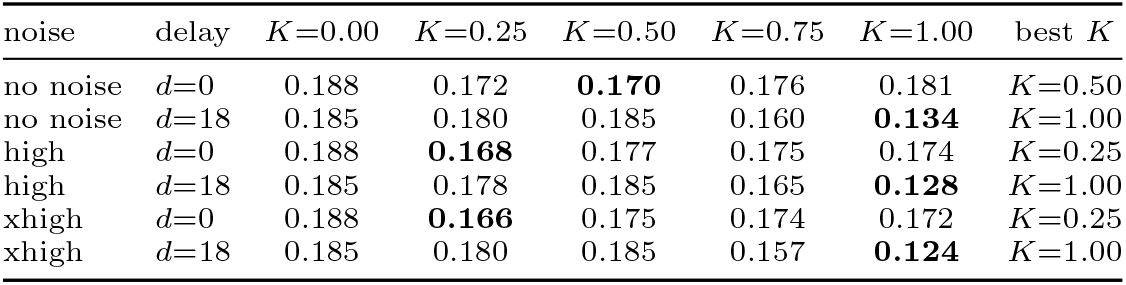
Per-cell swept fixed-*K* best *K* and corresponding cell-mean min-tip error (m), *n*=200 episodes per cell, stabilized endpoint probe, *H*=8 forward model, below-shoulder target set.

Table 1 reports the exact per-cell best *K* and min-tip mean. The diagonal trend *K*^⋆^ = 0.50 → 0.25 at delay-0 as noise rises, and the saturation at *K*^⋆^ = 1.0 across all delay-18 cells, are statistically robust under the paired bootstrap of Sec. 4.3 (every *K*=0 vs best-*K* contrast excludes zero at the 95% percentile level).

### 4.3 Controller-health diagnostics reclassify *K*=0 as an ablation (C2)

The *K* = 0 condition is valuable precisely because it reveals a failure mode: autoregressive forward-model drift generates controller commands that are hard to realize in muscle space. We log two controller-health diagnostics during every closed-loop episode — the NNLS residual ∥*M* ^⊤^*a* −*u*_joint_∥_2_ from Eq. (17), which measures how far the requested joint command is from anything achievable by non-negative muscle activations, and the saturation fraction of *a* at 0.95 — and a paired 95% bootstrap contrast against the per-cell best fixed *K*.

Figure 3A shows the paired 95% bootstrap CI of min-tip difference, *K*=0 minus best *K*, per cell. The CI excludes zero in every cell; the mean penalty is 1.9–2.2 cm at delay-0 cells and 5.2–6.1 cm at delay-18 cells. Panel B shows the NNLS residual mean heatmap: *K* = 0 runs sit at 92–96 across the grid, while *K >* 0 runs sit between 11 and 49. The same trend holds in representative single-episode traces (Appendix C): under *K* = 0, the forward-model state vector drifts by meters over the 600-step episode, the joint torque command requested by the task-space PD diverges, and NNLS returns large residuals because no non-negative muscle activation can realize that drifting command.

We therefore report *K* = 0 as a diagnostic ablation in all subsequent comparisons, not as the closed-loop oracle. The *h*=50 open-loop validation that selected *H*=8 is insufficient to certify *K* = 0 for closed-loop deployment, because the deployment horizon is 600 steps under a controller-induced action distribution that the forward model never trained on; we leave target-field clamping and controller-induced action augmentation as future work (see Sec. 5).

### 4.4 Outcome-trained reliability adaptation improves delayed reaches (C3)

We now ask whether agent-available signals — per-episode innovation history and per-episode min-tip outcome — can shift the deployed correction gain toward the swept oracle. We compare four estimators on top of the same *H*=8 forward model, the same stabilized endpoint probe, and the same 200-ep/cell deployment:

- **Default reliability** (*β*_0,*f*_ = 0, *β*_1,*f*_ = 0.5, all fields), an EMA-based reliability-to-gain logistic with no training signal.
- **Single-cell SPSA** on the no-noise, delay-18 cell, deployed on that one cell only.
- **Global SPSA** *β*, a single *β* trained across all six cells via outcome-only SPSA (100 iter, 12 samples per side; Sec. 3.5).
- **Feature-conditioned** *β*, a 30-parameter linear adapter that maps z-scored log innovation statistics to per-field *β* on top of the global SPSA base (Sec. 3.6).

For reference we also deploy a **per-cell SPSA diagnostic** (100 iter, 10 samples per side, one *β* per cell, deployed on its own training cell at *n*=200); this is an upper bound on what the reliability-to-gain logistic can achieve with within-cell training and is not an agent-realistic policy.

Figure 4 compares the default, global, feature-conditioned, and per-cell deployments against the swept fixed-*K* oracle, split by delay. At delay-18 cells, outcome-driven adaptation is unambiguous: global SPSA improves on default reliability by − 1.85 cm in the no-noise, delay-18 cell, − 2.18 cm at (high, *d*=18), and − 2.46 cm at (xhigh, *d*=18), and the paired 95% bootstrap CI excludes zero in every delay-18 cell (Table 2). The single-cell SPSA on the no-noise, delay-18 cell reduces min-tip by 2.50 cm compared with the default on that same cell, consistent with the global SPSA result. At delay-0 cells, neither global SPSA nor feature-conditioned *β* improves on default (Δ within ±1 cm; paired CI covers zero in every delay-0 cell). This split aligns directly with the delay-dependent oracle of C1: in cells where the swept oracle selects *K* = 1, outcome-driven adaptation can find that direction; in cells where the oracle is intermediate, the same parameterization has little to gain over the default mid-*K* regime.

**Table 2.** Key paired-bootstrap 95% percentile CIs on min-tip mean differences, in meters. Each contrast is computed as the first method named minus the second. All deploys at *n*=200 episodes/cell, paired by episode index, 10,000 resamples. Asterisk marks intervals that exclude zero. The complete CSV is regenerated by the reproduction pipeline described in Appendix D.

| contrast | cell | $\Delta$ | CI low | CI high |
| --- | --- | --- | --- | --- |
| global – default | no noise, $d=18$ | −0.0185 | −0.0241 | −0.0131 * |
| | high, $d=18$ | −0.0218 | −0.0286 | −0.0154 * |
| | xhigh, $d=18$ | −0.0246 | −0.0325 | −0.0168 * |
| | no noise, $d=0$ | −0.0030 | −0.0083 | +0.0020 |
| | high, $d=0$ | +0.0022 | −0.0057 | +0.0095 |
| | xhigh, $d=0$ | +0.0054 | −0.0015 | +0.0125 |
| feature-conditioned – best $K$ | no noise, $d=18$ | +0.0162 | +0.0097 | +0.0229 * |
| | high, $d=18$ | +0.0166 | +0.0092 | +0.0241 * |
| | xhigh, $d=18$ | +0.0174 | +0.0115 | +0.0235 * |
| $K=0$ – best $K$ | no noise, $d=18$ | +0.0516 | +0.0441 | +0.0595 * |
| | high, $d=18$ | +0.0570 | +0.0489 | +0.0652 * |
| | xhigh, $d=18$ | +0.0613 | +0.0542 | +0.0685 * |
| | no noise, $d=0$ | +0.0187 | +0.0118 | +0.0257 * |
| | high, $d=0$ | +0.0201 | +0.0131 | +0.0274 * |
| | xhigh, $d=0$ | +0.0218 | +0.0144 | +0.0290 * |

The full paired-bootstrap contrast table is generated by the reproduction pipeline in Appendix D.

### 4.5 The remaining gap to the swept oracle (C4)

The feature-conditioned *β* adapter does not reach the swept fixed-*K* oracle. Across the three delay-18 cells, feature-conditioned *β* minus best *K* is +1.6–1.7 cm, and the paired 95% CI excludes zero in every case (Table 2). The per-cell SPSA diagnostic, which is the strongest within-cell reliability-to-gain learner this parameterization allows, behaves similarly: in 5*/*6 cells it is statistically indistinguishable from feature-conditioned *β*, and in (xhigh, *d*=0) it improves on feature-conditioned *β* by 1.2 cm. Both estimators remain 1.4–1.8 cm above the swept oracle at every delay-18 cell.

Figure 5 explains *what* the adaptive rule actually did. Under the default (*β*_0,*f*_, *β*_1,*f*_)=(0, 0.5) rule, the realized correction gain (second-half episode mean) sits between 0.39 and 0.92 depending on the field. Under global SPSA, the `qpos` gain rises to 0.88–0.98 across cells while `qvel` is suppressed to 0.34. Feature-conditioned *β* shifts the same `qpos` field further toward 0.90–0.98 on a percell basis. The adaptive rule therefore reshapes field-wise trust, not merely scalar *K*; even so, the residual gap to the swept oracle persists, indicating that the reliability-to-gain logistic, the cell-level innovation features it conditions on, and the fixed-lag observer dynamics jointly bound the achievable adaptation under agent-available signals.

The adaptive deployment summary in Table 3 consolidates the headline cell-mean min-tip across all five estimators plus the swept oracle and the *K* = 0 diagnostic.

**Table 3.** Adaptive deployment summary, cell-mean min-tip error (m), *n*=200 episodes per cell. Boldface marks the smallest min-tip per row among the four adaptive estimators (default, global *β*, feature-conditioned *β*, per-cell *β*). The single-cell *β* diagnostic is reported only for the no-noise, delay-18 cell on which it was trained. *K*=0 is shown for diagnostic comparison and is not the closed-loop oracle (Sec. 4.3).

| noise | delay | default | global | feature-cond. | per-cell | single-cell | $K=0$ | best $K$ |
| --- | --- | --- | --- | --- | --- | --- | --- | --- |
| no noise | $d=0$ | 0.179 | 0.176 | **0.173** | 0.173 | – | 0.188 | 0.170 |
| no noise | $d=18$ | 0.173 | 0.155 | 0.150 | **0.148** | 0.148 | 0.185 | 0.134 |
| high | $d=0$ | **0.174** | 0.177 | 0.175 | 0.176 | – | 0.188 | 0.168 |
| high | $d=18$ | 0.174 | 0.152 | **0.145** | 0.145 | – | 0.185 | 0.128 |
| xhigh | $d=0$ | 0.169 | 0.175 | 0.173 | **0.161** | – | 0.188 | 0.166 |
| xhigh | $d=18$ | 0.173 | 0.148 | **0.141** | 0.142 | – | 0.185 | 0.124 |

## 5 Discussion

### Delay changes the role of sensory prediction error

The C1 fixed-*K* sweep separates two regimes that are hidden if *K* = 0 is treated as a usable closed-loop oracle rather than as a forward-model-only diagnostic. With no observation delay, the swept oracle sits at intermediate *K*∈ {0.25, 0.50}: the predictive observer benefits from balancing its forward-model prediction against the current sensor reading, and increasing observation noise pushes that optimum toward prediction, as expected from reliability-weighted prediction-error correction. With an 18-step delay, the swept oracle collapses to *K* = 1 in every noise condition: the forward-model prediction at the current step is itself stale (the observer’s input is delayed by the same amount as the observation), so observation-heavy correction wins even at the high-noise level. We describe this only as a qualitative reliability-weighted correction structure; the closed loop here is not optimal control, and the observer does not propagate covariances. *K* = 0 **is a forward-model-only diagnostic**. Under the stabilized endpoint probe, *K* = 0 is systematically worse than the per-cell best fixed *K* in every cell, with paired 95% CIs that exclude zero; the penalty is 5.2–6.1 cm at delay-18 cells. The NNLS controller residual at *K* = 0 is several times larger than at *K >* 0, indicating that the forward-model prediction received by the controller is not realizable as non-negative muscle activations. This is a deployment-distribution gap. The open-loop *h* = 50 MSE validation selected *H*=8 from training data generated by random, low-amplitude, and hold controllers; the closed-loop deployment runs 600 steps under a controller-induced action distribution that the forward model was never trained on, and the autoregressive rollout drifts to non-physical states. The implications for future work are concrete: the target field should be invariant or clamped rather than predicted autoregressively; training should include controller-induced action distributions if the model is used closed-loop at low *K*; and *h* = 600 stability should be evaluated under the same action distribution used at deployment. We retain *K* = 0 on the *K*-grid as a diagnostic to expose this failure mode, not as a candidate oracle.

### What outcome-trained reliability adaptation can and cannot do

The C3 result is delay-dependent in a way that follows directly from C1. In delay-18 cells, where the swept oracle is *K* = 1, outcome-driven SPSA on a single global *β* moves the realized correction toward observation-heavy correction; the field-wise breakdown (Fig. 5) shows the `qpos` gain rising from default ∼ 0.72 to ∼ 0.89 while `qvel` is suppressed. In delay-0 cells, where the swept oracle is intermediate, the same adaptation yields no improvement over the default mid-*K* regime. None of these results imply that the reliability-adaptive observer has *learned the optimal Kalman gain*: it has learned a *β* that the closed-loop outcome prefers, on a controller that is a deliberately stabilized probe, not a biological motor planner.

### The remaining adaptation gap

Even feature-conditioned *β*, which nearly matches the per-cell trained diagnostic in 5*/*6 cells, remains 1.4–1.8 cm worse than the swept fixed-*K* oracle at every delay-18 cell (Table 2). Four plausible contributors: (i) the field-wise reliability-to-gain logistic *K*_*f*_ = *σ*(*β*_0,*f*_ + *β*_1,*f*_ log *r*_*f*_) is a constrained parameterization that may not span the swept fixed-*K* solution; (ii) the 10-dimensional cell-level innovation summary used by the adapter may not encode the cell-specific structure that determines the best fixed *K*; (iii) the fixed-lag observer’s recursive dynamics differ from a scalar *K* applied to a clean innovation, so even an oracle *β* would not necessarily reproduce the swept-*K* outcome; (iv) controller and estimator are coupled, and the residual gap may reflect controller response to imperfect state estimates rather than estimation error itself. We do not separate these factors here; that decomposition is left to future work.

### Scope and limitations

(a) The main task is restricted to IK-reachable below-shoulder targets. The constraint is anatomically motivated (tabletop-like reaching) and avoids mechanically degenerate above-shoulder IK edge solutions, but it is not a substitute for evaluation on a freely varying set of targets. (b) The closed-loop controller is a stabilized probe, not a model of biological motor planning. The NNLS routing and virtual target ramp were introduced specifically to avoid coupled failure modes among the controller, IK solver, and forward model; we do not claim they are how the nervous system solves the same problem. (c) All adaptive results use a single *H*=8 main model, a single SPSA random seed, and a single *β* parameterization. We do not vary the forward-model architecture or supervision horizon in this study. (d) The *K* = 0 closed-loop instability is not necessarily a property of any forward-model-only observer; it is a property of this *H*=8 residual MLP trained on random, low-amplitude, and hold trajectories, deployed for 600 steps under a controller-induced action distribution. Forward models with target-field clamping or deployment-distribution-aware training could change the picture. (e) Closed-loop estimator benchmarks should carry controller-health metrics. The artifactual *K* = 0 optimum we initially observed would not have been detectable from min-tip alone; it was detectable from NNLS residual and field-wise realized *K*.

## 6 Conclusion

We have presented a stabilized closed-loop benchmark for a forward-model-based predictive state observer on a 34-muscle musculoskeletal reaching task, together with an analysis that separates four phenomena previously conflated by an earlier controller diagnostic.

First, the swept fixed-*K* closed-loop oracle is delay-dependent (Sec. 4.2): at delay-0 cells, the best fixed *K* is intermediate (*K* = 0.50 in the no-noise cell and *K* = 0.25 at higher noise), while at delay-18 cells the best fixed *K* collapses to *K* = 1.0 across all noise levels. Second, the forward-model-only *K* = 0 condition is systematically worse than the per-cell best fixed *K* in every cell (paired 95% CIs exclude zero; mean penalty 1.9–6.1 cm) and shows large NNLS controller residuals caused by long-horizon autoregressive forward-model drift (Sec. 4.3); we therefore report *K* = 0 as a diagnostic forward-model-only ablation, not as a closed-loop oracle. Third, outcome-trained reliability adaptation improves the delayed regime: a single global *β* trained via outcome-only SPSA reduces min-tip error by 1.85–2.46 cm versus default reliability at every delay-18 cell (all paired 95% CIs exclude zero), while delay-0 cells remain statistically neutral (Sec. 4.4); this delay-dependent benefit follows directly from the swept-*K* structure. Fourth, a feature-conditioned *β* adapter nearly matches a per-cell trained reliability-to-gain diagnostic in 5*/*6 cells, but both adapters remain 1.4–1.8 cm worse than the swept fixed-*K* oracle at every delay-18 cell (Sec. 4.5). The remaining gap indicates that the reliability-to-gain logistic, the cell-level innovation features it conditions on, and the fixed-lag observer dynamics jointly bound what agent-available adaptation can deliver under this controller.

The takeaway is therefore methodological as well as substantive: closed-loop state-estimation benchmarks need controller-health and long-horizon diagnostics to distinguish a state-estimation finding from artifacts caused by coupling among the controller, IK solver, and forward model; under a stabilized probe controller, agent-available reliability and outcome signals can shift the deployed correction gain in the right direction when the oracle is observation-heavy, but cannot eliminate the remaining oracle gap with this parameterization. Future work should train forward models on controller-induced action distributions and test *β* parameterizations richer than the field-wise logistic.

## Acknowledgments

The author thanks the MyoSuite maintainers for releasing the musculoskeletal benchmark used in this study.

## Statements and Declarations

### Ethics approval

Not applicable. This study uses only the publicly available MyoSuite musculoskeletal simulation environment in MuJoCo; no human, animal, or clinical data are involved.

### Consent to participate and consent for publication

Not applicable (no human subjects).

### Data availability

All main-analysis data are generated from the MyoSuite `myoArmReachFixed-v0` environment Caggiano et al. (2022); historical diagnostics retained in the repository also include `myoArmReachRandom-v0`. Generation scripts and configurations are included with the code release (Appendix D).

### Code availability

The full reproduction pipeline (dataset generation, forward-model training, closed-loop evaluation, across-trial *β* adaptation, feature-conditioned *β* training, and figure generation) is released as open-source software at https://github.com/jkoba0512/myoarm-forward-state-estimation; an archived release DOI will be added upon acceptance. Frozen run identifiers required to regenerate every numerical claim and figure are listed in Appendix D.

### Author contributions

Jun Kobayashi is the sole author and conducted all aspects of the work: conceptualization, methodology, software, experiments, analysis, and manuscript preparation.

## Appendix A Implementation Details

Table A1 lists every hyperparameter used in the paper. All training runs are reproducible from `configs/` in the source repository; estimator evaluation grids are listed verbatim in `configs/estimators/` and `configs/closed loop/`.

## Appendix B Forward-Model Rollout Metrics and *h*=600 Stability

### Open-loop rollout error

Table B2 reports horizon-wise prediction error for the main forward-model variants (single-step-supervised residual MLP on the 77,292-transition reachable-target dataset; same architecture and training set, varied only in the rollout horizon *H*). The *H*=8 main model is the one used throughout the closed-loop results; an undertrained *H*=8 variant is retained for diagnostic contrast.

### h=600 zero-action stability

Because the closed-loop episode is 600 control steps long, we additionally evaluate a zero-action autoregressive rollout from `env.reset()` on every forward-model variant (Table B3). The field-wise stability screen is |*q*| _max_ ≤ 5 rad, 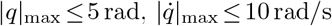, tip drift on the order of the anatomical workspace (∼ 1 m), and no NaN or Inf values. Target drift is reported as a diagnostic field only — all four models drift in *p*_tgt_ because the residual-MLP predicts the full state vector, and the target field is included in that state; in a future iteration the target field should be clamped to their initial value or removed from the predicted state. In closed-loop deployments, the target channel is observed and corrected with fixed gain 1.0; the reported target drift therefore diagnoses only the unconstrained open-loop rollout, not the controller-facing target estimate.

### Closed-loop K=0 instability

Under *K*=0 the same *H*=8 main model is driven by a controller-induced action sequence (rather than zero action) and the rollout diverges much more aggressively (e.g. |*q*|_max_=11.8 rad,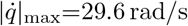, tip drift 8.76 m at the same 600-step horizon). The gap between the zero-action and the closed-loop *K*=0 rollouts is the deployment-distribution gap discussed in Sec. 4.3; it motivates including controller-induced action distributions in future forward-model training.

**Table A1.** Hyperparameters used across the paper.

| Component | Hyperparameter | Value |
| --- | --- | --- |
| Forward model $f_\theta$ | state dimension | 83 |
|  | action dimension | 34 |
| | hidden layers | $2 \times 256$ |
|  | activation | ReLU, LayerNorm |
| | output | residual ($\Delta x$) |
|  | parameters | 118,355 |
| Forward training | optimizer | Adam Kingma and Ba (2015) |
| | learning rate | $1 \times 10^{-3}$ |
|  | weight decay | 0 |
|  | batch size | 256 |
|  | epochs | 200 (early stop patience 20) |
|  | validation split | every 5th source episode |
| Multi-step loss | rollout horizon $H$ | 1, 4, 8 |
| Stabilized endpoint probe | $K_p, K_d$ | 30, 3 |
| | action scale $\alpha$ | 5 |
|  | ramp horizon | 300 steps |
| | muscle routing | non-negative least squares, then clip to $[0, 1]$ |
| IK solver | method | damped Jacobian pseudoinverse |
|  | max iterations / tolerance | 200 / 0.01 m |
| | damping $\lambda^2$ | 0.1 |
| Eval grid (open-loop) | noise levels | none (no noise), low, medium, high, vhigh, xhigh |
|  | delays (steps) | 0, 6, 18, 36 |
| | $K$ values | 0.0, 0.25, 0.5, 0.75, 1.0 |
| Eval grid (closed-loop) | noise levels (focused) | none (no noise), high, xhigh |
|  | delays (focused) | 0, 18 |
| Closed-loop deploys | episodes per cell | 200 |

**Table B2.** Forward-model rollout MSE and tip prediction error on held-out validation episodes for the reachable-target dataset (lower is better).

| Model | MSE |  |  | Tip err. (m) |  |  |
| --- | --- | --- | --- | --- | --- | --- |
| | $h=1$ | $h=10$ | $h=50$ | $h=1$ | $h=10$ | $h=50$ |
| $H=1$ (single-step) | 0.0035 | 0.0089 | 0.0366 | 0.0040 | 0.0157 | 0.0546 |
| $H=4$ multi-step | 0.0037 | 0.0060 | 0.0115 | 0.0064 | 0.0206 | 0.0538 |
| $H=8$ multi-step (main) | 0.0038 | 0.0057 | 0.0084 | 0.0071 | 0.0162 | 0.0343 |
| undertrained $H=8$ | 0.0055 | 0.0082 | 0.0110 | 0.0134 | 0.0267 | 0.0517 |

**Table B3.** Zero-action *h*=600 autoregressive stability per forward-model variant. The two *H*=8 variants remain bounded in 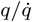 and keep tip drift near the anatomical-workspace scale (target drift treated as a diagnostic field). *H*=1 and *H*=4 diverge to non-physical states.

| Model | $\vert q\vert _{\max}$ (rad) | $\vert \dot{q}\vert _{\max}$ (rad/s) | tip drift (m) | tgt drift (m) | status |
| --- | --- | --- | --- | --- | --- |
| $H=1$ | 15.6 | 108.0 | 7.76 | 0.28 | diverges |
| $H=4$ | 42.4 | 217.4 | 12.6 | 10.2 | diverges |
| $H=8$ (main) | 1.6 | 3.9 | 1.03 | 0.54 | bounded |
| undertrained $H=8$ | 1.5 | 3.4 | 0.43 | 0.96 | bounded |

## Appendix C Below-Shoulder Target Set and Historical Controller Diagnostic

The main target set is generated by uniformly sampling the environment’s task-space reach bounding box, applying a damped-Jacobian IK acceptance test (tolerance 1 cm), and rejecting candidates above the neutral shoulder height (*z* = 1.393 m). This shoulder-height criterion is anatomical, not a post-hoc performance filter: it removes mechanically degenerate above-shoulder reaches that the IK solver can satisfy only by pinning several shoulder/scapula joints to range limits. The acceptance rate is 55.9% for *n*=200 accepted targets; after the shoulder-height filter, the IK tolerance test rejects 0% of candidates.

The change to the stabilized endpoint controller was motivated by an earlier diagnostic that used a one-shot IK pre-solve, a joint-space PD controller, a ReLU-then-moment-arm muscle map, and an unrestricted target set. That diagnostic produced a misleading apparent optimum at *K* = 0: high-reaching targets pinned joints, the muscle map could not realize the resulting joint command, and long-horizon forward-model autoregression drifted to non-physical states. The stabilized endpoint probe (Sec. 3.7) and the below-shoulder target set remove these failure modes; the historical diagnostic is not used as a main result.

## Appendix D Software, Hardware, and Reproducibility

### Software stack

Python 3.11, PyTorch configured for CPU execution, MuJoCo 3.x, Gymnasium 0.29, MyoSuite 2.12.2, NumPy, SciPy, pandas, and matplotlib. Project-level dependencies are pinned in `pyproject.toml.`

### Hardware

All experiments were run on a single CPU workstation (no GPU required). Forward-model training takes ∼ 30 min per model. A small fixed-*K* diagnostic sweep of 300 episodes (5 values of *K* × 6 cells ×10 episodes) takes ∼ 10 min; the main 200-episode-per-cell deployment tables take proportionally longer. Per-cell SPSA training (100 iterations, *S* = 10, 20 episodes per iteration) takes ∼1 hour per cell when run alone; the 200-iteration feature-conditioned *β* training (Sec. 3.6) takes ∼ 3.5 hours.

### Main reproduction path

The repository contains executable configuration files for each training, evaluation, and plotting step. The main numerical tables and figures are regenerated from the fixed-gain closed-loop sweep config, the adaptive-deployment configs, the paired-bootstrap wrapper, and the paper-figure builder listed in the released source tree. The forward-model and SPSA training commands are listed in `configs/models/ and configs/train/;` each run writes its config and Git commit hash to `runs/`.

### Historical reproduction

Earlier joint-PD diagnostics are retained in the repository for auditability but are excluded from the main analysis because they used the unrestricted target set and controller pipeline described in Appendix C. They should not be used to reproduce the stabilized-controller results.

### Repository layout

~~~
src/myoarm_fse/
 envs/ MyoSuite wrappers, observation pipelines
 models/ ForwardMLP, training loops
 estimators/ FixedGainKalman, ReliabilityAdaptive, Learned
 controllers/ JointSpacePD, EndpointError
 controllers/ BC kept for historical runs
 evaluation/ closed_loop.py (single episode rollout)
scripts/ training, evaluation, and plotting entries
configs/
 models/ forward-model training configs
 closed_loop/ closed-loop eval configs
 train/ SPSA and per-cell SPSA configs
 train/ feature-conditioned beta configs
runs/ local outputs; large artifacts not versioned
figures/ committed PDF/PNG of paper figures
paper/ LaTeX source + bibliography
~~~

### Tests

Unit tests cover the per-step observer math, fixed-lag delay handling, and the deterministic build of every estimator. A MyoSuite-marked smoke suite verifies that one closed-loop episode runs end-to-end with each controller-estimator combination. The reproduction checklist uses:

~~~
uv run pytest # default fast tests
uv run pytest -m myosuite # closed-loop integration smoke
~~~

### Run artifacts

The repository includes configs that, when run, write reproducible output directories under `runs/`. Each closed-loop or training run records its resolved config and a Git commit hash in `config.json`, so re-running with the same seed and commit reproduces the reported seeded outputs. Frozen run identifiers are embedded in the configs and wrapper scripts listed above, and are kept there rather than repeated in the manuscript text.

## References

Bernstein, N.A.: The Coordination and Regulation of Movements. Pergamon Press, Oxford (1967)

Wolpert, D.M., Ghahramani, Z.: Computational principles of movement neuroscience. Nature Neuroscience 3(11), 1212–1217 (2000) 10.1038/81497

Kalman, R.E.: A new approach to linear filtering and prediction problems. Transactions of the ASME–Journal of Basic Engineering 82(1), 35–45 (1960) 10.1115/1.3662552

Mehra, R.K.: On the identification of variances and adaptive Kalman filtering. IEEE Transactions on Automatic Control 15(2), 175–184 (1970) 10.1109/TAC.1970.1099422

Anderson, B.D.O., Moore, J.B.: Optimal Filtering. Prentice-Hall, Englewood Cliffs, NJ (1979). Republished by Dover, 2005.

Spall, J.C.: Multivariate stochastic approximation using a simultaneous perturbation gradient approximation. IEEE Transactions on Automatic Control 37(3), 332–341 (1992) 10.1109/9.119632

Caggiano, V., Wang, H., Durandau, G., Sartori, M., Kumar, V.: MyoSuite – a contact-rich simulation suite for musculoskeletal motor control. In: Proc. 4th Annual Learning for Dynamics and Control Conference (L4DC). Proceedings of Machine Learning Research, vol. 168, pp. 492–507. PMLR, PMLR (2022). https://proceedings.mlr.press/v168/caggiano22a.html

Todorov, E., Erez, T., Tassa, Y.: MuJoCo: A physics engine for model-based control. In: Proc. IEEE/RSJ Int. Conf. Intelligent Robots and Systems (IROS), pp. 5026–5033 (2012). 10.1109/IROS.2012.6386109

Rauch, H.E., Tung, F., Striebel, C.T.: Maximum likelihood estimates of linear dynamic systems. AIAA Journal 3(8), 1445–1450 (1965) 10.2514/3.3166

Hasani, R., Lechner, M., Amini, A., Rus, D., Grosu, R.: Liquid time-constant networks. In: Proc. AAAI Conf. Artificial Intelligence, vol. 35, pp. 7657–7666 (2021). https://ojs.aaai.org/index.php/AAAI/article/view/16936

Hasani, R., Lechner, M., Amini, A., Liebenwein, L., Ray, A., Tschaikowski, M., Teschl, G., Rus, D.: Closed-form continuous-time neural networks. Nature Machine Intelligence 4(11), 992–1003 (2022) 10.1038/s42256-022-00556-7

Hafner, D., Lillicrap, T., Ba, J., Norouzi, M.: Dream to control: Learning behaviors by latent imagination. In: Proc. Int. Conf. Learning Representations (ICLR) (2020). https://arxiv.org/abs/1912.01603

Janner, M., Fu, J., Zhang, M., Levine, S.: When to trust your model: Model-based policy optimization. In: Advances in Neural Information Processing Systems, vol. 32 (2019). https://proceedings.neurips.cc/paper/2019/hash/5faf461eff3099671ad63c6f3f094f7f-Abstract.html

Pomerleau, D.A.: ALVINN: An autonomous land vehicle in a neural network. In: Advances in Neural Information Processing Systems, vol. 1. Morgan Kaufmann, San Mateo, CA (1988). https://proceedings.neurips.cc/paper/1988/hash/812b4ba287f5ee0bc9d43bbf5bbe87fb-Abstract.html

Ross, S., Gordon, G.J., Bagnell, J.A.: A reduction of imitation learning and structured prediction to no-regret online learning. In: Proc. 14th Int. Conf. Artificial Intelligence and Statistics (AISTATS). Proceedings of Machine Learning Research, vol. 15, pp. 627–635. PMLR, Fort Lauderdale, FL (2011). https://proceedings.mlr.press/v15/ross11a.html

Kingma, D.P., Ba, J.: Adam: A method for stochastic optimization. In: Proc. 3rd Int. Conf. Learning Representations (ICLR) (2015). https://arxiv.org/abs/1412.6980

